# The RNA helicases DDX5 and DDX17 facilitate neural differentiation of human pluripotent stem cells NTERA2

**DOI:** 10.1101/2021.05.10.443309

**Authors:** Praewa Suthapot, Tiaojiang Xiao, Gary Felsenfeld, Suradej Hongeng, Patompon Wongtrakoongate

## Abstract

Understanding human neurogenesis is critical toward regenerative medicine for neurodegeneration. However, little is known how neural differentiation is regulated by RNA helicases, which comprise a diverse class of RNA remodeling enzymes. We show here that expression of the DEAD box-containing RNA helicases DDX5 and DDX17 is abundant throughout retinoic acid-induced neural differentiation of the human pluripotent stem cell (hPSC) line NTERA2, and is mostly localized within the nucleus. Using ChIP-seq, we identify that the two RNA helicases occupy chromatin genome-wide at regions associated with neurogenesis- and differentiation-related genes in both hPSCs and their neural derivatives. Further, RNA-seq analyses indicate both DDX5 and DDX17 are mutually required for controlling transcriptional expression of these genes. We show that the two RNA helicases are not important for maintenance of stem cell state of hPSCs. In contrast, they facilitate early neural differentiation of hPSCs, generation of neurospheres from the stem cells, and expression of key neurogenic transcription factors during neural differentiation. Importantly, DDX5 and DDX17 are important for differentiation of hPSCs toward NESTIN- and TUBB3-positive cells, which represent neural progenitors and mature neurons. Collectively, our findings suggest the role of DDX5 and DDX17 in transcriptional regulation of genes involved in neurogenesis, and hence in neural differentiation of hPSCs.

## Introduction

Embryonic neurogenesis initiates approximately at gestational days 9–9.5 and 24-28 in mice and men, respectively (DeSesso, Scialli et al. 1999, Rice and Barone 2000), by the commitment of post-implantation epiblasts into embryonic neural precursors, which subsequently generate neural derivatives including neurons, astrocytes, and oligodendrocytes (Zhang, Wernig et al. 2001, Götz and Huttner 2005). In embryonic and fetal brains, neurogenesis is found mostly at the dentate gyrus of the hippocampus and the subventricular zone of the lateral ventricles. Dysregulation of neurogenesis has been shown to associate with various neurodevelopmental disorders and neurodegenerative diseases such as Rett syndrome (Shahbazian and Zoghbi 2002, Tudor, Akbarian et al. 2002), schizophrenia (Reif, Fritzen et al. 2006, Duan, Chang et al. 2007) and Alzheimer disease (Jin, Peel et al. 2004, Verret, Jankowsky et al. 2007). Hence, it is crucial to dissect mechanisms controlling neurogenesis at molecular and cellular levels. In addition, although progresses have been made to understand mouse embryonic neurogenesis, little is known for the human counterpart.

In human pluripotent stem cells (hPSCs), embryonic neurogenesis can be initiated by activation of the neurogenic transcription factor PAX6 (Götz, Stoykova et al. 1998, Zhang, Huang et al. 2010), which then further controls the expression program of other key neurogenic transcription factors including SOX1 (Genethliou, Panayiotou et al. 2009, Suter, Tirefort et al. 2009), SOX2 (Kondoh, Uchikawa et al. 2004, Wen, Hu et al. 2008), SOX21 (Sansom, Griffiths et al. 2009), ASCL1 (Sansom, Griffiths et al. 2009) and NEUROG2 (Scardigli, Baumer et al. 2003, Bel-Vialar, Medevielle et al. 2007). We have previously reported that an ectopic expression of PAX6 in the hPSC line NTERA2 led to a massive generation of embryonic neural precursors (Sutiwisesak, Kitiyanant et al. 2014). Apart from DNA-binding transcription factors, neurogenesis is also tightly controlled by RNA metabolisms (Loya, Van Vactor et al. 2010, Di Liegro, Schiera et al. 2014) such as RNA-binding proteins and RNA helicases. The latter has been shown to control multi-step gene expression including chromatin biology, transcription, post-transcription, and translation (Abdelhaleem 2009, Fuller-Pace 2013, Robert and Pelletier 2013). For example, the RNA helicase DDX6 has been proposed to induce neurogenesis via Let-7a-mediated post-transcriptional control (Nicklas, Okawa et al. 2015). Further, translational initiation of Mical mRNA has been found to be tightly controlled by the DEAD-box RNA helicase eIF4A, which is indispensable for dendrite pruning of *Drosophila* sensory neurons (Rode, Ohm et al. 2018). Since many RNA helicases have been identified for their physical property to occupy chromatin and to exert their effect via transcriptional co-regulation (Tanaka, Okamoto et al. 2009, Johansson, Marie et al. 2020, Pérez-Calero, Bayona-Feliu et al. 2020), it is also yet to be elucidated whether these RNA helicases regulate neurogenesis by genome-wide transcriptional control.

Among the RNA helicase family, DDX5 and DDX17 are orthologous DEAD-box RNA helicases which share higher than 90% sequence similarity across the central core region suggesting their conserved molecular functions. Nonetheless, the differences at amino- and carboxy-terminal amino acid sequences may provide additional and specific function for these individual RNA helicases (Lamm, Nicol et al. 1996, Fuller-pace 2006, Fuller-pace and Ali 2008). The RNA helicases DDX5 and DDX17 play a role in the regulation of transcriptional initiation by acting as a co-activator or a corepressor involving cell growth and differentiation. For instance, DDX5 and DDX17 have been reported to form a complex with beta-catenin and to control epithelial-mesenchymal transition (Yang, Lin et al. 2006, Shin, Rossow et al. 2007). They can also interact with the Notch intracellular domain and facilitate the expression of Notch target genes including *preTCRα, Hes1, and CD25* (Jung, Mittler et al. 2013). In addition, we have shown that DDX5 can participate in chromatin long-range interaction mediated by the chromatin architectural protein CTCF (Yao, Brick et al. 2010). We and others have also demonstrated that DDX5 occupies chromatin genome-wide in hPSCs (Wongtrakoongate, Riddick et al. 2015), HeLa cells (Yao, Brick et al. 2010), spermatogonia (Legrand, Chan et al. 2019), and breast cancer (He, Song et al. 2019) indicating its genome regulation in these cell types, and possibly in neurogenesis.

In the present study, we aimed to characterize chromatin occupancy of the RNA helicases DDX5 and DDX17 in the hPSCs NTERA2 and their neural derivatives. We find that both DDX5 and DDX17 are mutually required for transcriptional activation of neural differentiation-associated genes. Even though the two RNA helicases are not important for maintenance of the stem cell state of hPSCs, they are nonetheless critical for neural differentiation of hPSCs. Thus, our works uncovered the transcriptional regulation of DDX5 and DDX17 in human neurogenesis.

## Materials and methods

### Cell culture and differentiation of NTERA2 cells

The human embryonal carcinoma stem cells NTERA-2 cl.D1 (NT2/D1) was provided by Prof. Peter W. Andrews, University of Sheffield, UK. The cells were maintained in high glucose Dulbecco’s modified Eagle’s medium (HyClone, SH30243.02) supplemented with 10% EmbryoMax fetal bovine serum (Merck, ES-009-B) at 37°C with 5% CO2 and passaged every 3-4 days. For differentiation with retinoic acid, NTERA2 cells were trypsinized and plated at density 1×10^4^ cells/cm^2^ in culture medium containing 10 μM all-trans retinoic acid (Sigma, R2625), the fresh medium was replaced every four days.

For induction of neurosphere formation, cells were trypsinized with 0.05% Trypsin-EDTA (HyClone, SH30042.02) and seeded in a non-treated 48-well plate with a density of 1,000 cells/well. Cells were grown in DMEM/F12 (HyClone, SH30023.02) supplemented with 2 %v/v N21 (R&D system, AR008), 20 ng/mL EGF (R&D system, 236-EG), and 20 ng/mL bFGF (R&D system, 4114-TC) and cultivated under standard atmosphere (37°C, 5% CO2). Counting of neurosphere numbers was performed at day 8 post-treatment.

### RNA silencing

The ON-TARGETplus SMARTpool siRNA specific for DDX5 (L-003774-00-0010) and DDX17 (L-013450-01-0010) and the non-targeting control pool (D-001810-10-20) were purchased from Dharmacon. hPSCs and day 14 retinoic acid-induced neural derivatives were seeded at density of 3.3×10^4^ and 1×10^5^ cells/cm^2^, respectively, one day before siRNA transfection. Cells were transfected with 10 nM of siDDX5 and/or siDDX17 using Lipofectamine RNAiMAX Transfection Reagent (Invitrogen, #13778075). The non-targeting siRNA at 20 nM was used as a control transfection. After 60-hour post-transfection, cells were subsequently analyzed or induced to differentiate further as indicated.

### RNA extraction and quantitative realtime PCR

Total RNA was extracted using Genezol reagent (Geneaid Biotech, GZR100), and reversed transcribed by ReverTra Ace qPCR RT kit (Toyobo, FSQ-301-200) according to manufacturer’s instructions. Gene expression was determined by quantitative realtime PCR using SYBR Green Master Mix (PCR Biosystems, PB20.11-05) and normalized with the internal control GAPDH. Primer sequences are available upon request.

### Western blot and immunofluorescence

Cells were lysed with M-PER Mammalian Protein Extraction Reagent (Thermo Scientific, 78501) supplemented with protease inhibitor cocktail (Roche, 11836170001). The lysates were incubated for 10 minutes at room temperature and centrifuged at 14,000 rpm 4°C 15 minutes to remove debris. Aliquots of 30 μg of protein extracts were resolved on 10% SDS-PAGE before electrotransfer to a PVDF membrane. The membrane was subjected to western blot analysis with antibodies against proteins of interest as following; anti-DDX5 (Abcam, ab126730, 1:10,000 dilution), anti-DDX17 (Proteintech, 19910-1-AP, 1:3,000 dilution), anti-OCT4 (Invitrogen, MA1-104, 1:1,000 dilution) and anti-TUBB3 (Origene, TA500047, 1:3,000 dilution), anti-GAPDH conjugated peroxidase (Proteintech, HRP-60004, 1:30,000 dilution), goat anti-rabbit IgG conjugated peroxidase (Cell Signaling, #7074, 1:5,000 dilution), and horse anti-mouse IgG conjugated peroxidase (Cell Signaling, #7076, 1:5,000 dilution).

For detection of DDX5 and DDX17 expression and cell stage markers, undifferentiated NTERA2 and its neural derivative cultures were plated as single-cell into a 24-well plate for 1-2 days before conducting immunocytochemistry. Cells were fixed in 4% paraformaldehyde for 15 minutes at room temperature and permeabilized using 0.2% Triton-X-100 in PBS for another 15 minutes. Fixed cells were then blocked in 2% BSA in 0.1 % PBST for 1 hour and co-stained with primary antibodies against DDX5 or DDX17 and stage-specific markers with following dilutions; anti-DDX5 (Abcam, 1:300 dilution), anti-DDX17 (Proteintech, 1:150 dilution), anti-OCT4 (Invitrogen, 1:150 dilution), and anti-TUBB3 (Santa Cruz, sc-51670, 1:150 dilution). The primary antibodies were incubated overnight at 4°C. AlexaFluor 488 conjugated goat anti-mouse IgG (Cell Signaling, #4408) and AlexaFluor 594 conjugated goat anti-rabbit IgG (Cell Signaling, #8889) were applied to the samples at dilution of 1:300 with 1-hour incubation time.

For measuring neuron yield production, hPSC depleted for DDX5 and DDX17 were differentiated by retinoic acid, and continuously grown as a monolayer culture until day 16. The cells were dissociated by 0.05% Trypsin-EDTA and re-plated with dilution 1:5 in a 24-well plate a day before performing immunofluorescent staining assay. In parallel, neural derivative culture at day 14 was trypsinized and re-seeded at density 150,000 cells/cm^2^ in a 12-well plate prior to carry on RNA silencing. Cells were resumed growing in the retinoic acid-containing medium after 60 h and immunofluorescent staining was performed on day 32. On the analysis day, cells were fixed and stained according to the procedure described above using primary antibodies against NESTIN (Merck, ABD69, 1:300 dilution) and TUBB3 (Santa Cruz, 1:150 dilution). Cells were analyzed with a fluorescence microscope (Olympus, IX83ZDC) and image processing was done in the ImageJ program (NIH, USA).

### Chromatin Immunoprecipitation-sequencing

Chromatin immunoprecipitation followed by high-throughput DNA sequencing was performed in undifferentiated NTERA2 cells and retinoic acid-treated culture for 14 days. The cells were processed by formaldehyde fixation, nuclei isolation, and mechanical DNA fragmentation using a ChIP-IT Express kit (Active Motif, #53008) and Bioruptor (Diagenode) according to the protocol previously described (Wongtrakoongate, Riddick et al. 2015). An aliquot of 50 μg sheared chromatin was used per one IP reaction together with 3 μg antibody and the samples were incubated overnight at 4°C, 10% of chromatin solution was reserved as “input DNA”. The following antibodies were used for IP reaction: DDX5 (Bethyl Laboratories, A300-523A), DDX17 (Bethyl Laboratories, A300-509A), normal rabbit-IgG (Santa Cruz, sc-2027). The DNA fragments were purified by QIAquick PCR Purification Kit (Qiagen, #28104) subjected to library preparation according to the manufacturer’s instruction by using MicroPlex Library Preparation Kit (Diagenode). The ligated DNA (size 250–300 bp) was enriched by PCR before submitted to the NIDDK Genomic Core Facility for high-throughput sequencing using Illumina HiSeq2500. The sequencing data were aligned with the genome reference hg19/GRCh37 and exported into BAM file format. Sequencing data were submitted to GEO Datasets under accession number GSE174051.

The aligned reads were filtered with the SAMTools program to remove duplicates and peakcalling was performed in MACS version 1.4.2 with a threshold of p-value less than 10^-5^. The resulting BED files from each of biological triplicate samples were intersected and further analyzed by using the Bioconductor packages ChIPpeakAnno (Zhu, Gazin et al. 2010) and ChIPseeker (Yu, Wang et al. 2015), respectively

### RNA sequencing

Three replicates of control knockdown, individual knockdown of DDX5 or DDX17, and double knockdown treatment in hPSC and neural derivative cultures from 60-hour post-transfection were used for RNA library preparation, respectively. Total RNA was extracted from each knockdown sample using RNeasy Mini kit (Qiagen, #74104) and contaminated DNA was removed by using a TURBO DNA-free kit (Ambion, AM1907) following the manufacturer’s instructions. Samples were prepared using with 1 μg of total RNA and mRNA isolation step was done by using NEBNext poly(A) mRNA Magnetic Isolation kit (New England Biolabs, E7490S). The cDNA libraries were generated by the “dUTP” method for strand-specificity using NEBNext^®^ Ultra™ II Directional RNA Library Prep Kit for Illumina and NEBNext Multiplex Oligos for Illumina (New England Biolabs, E7760S, E7335). Library size (~350 bp) was checked by Bioanalyzer and the pool libraries from each biological replicate were sequenced on Illumina HiSeq 2500 in single-end-50 bp mode.

The raw sequencing reads in FASTQ format were trimmed with Trimmomatic (v.0.38) (Bolger, Lohse et al. 2014), aligned to human genome reference with HISAT2 (v.2.1.0+galaxy5, use a build-in genome: Human (hg19), reverse stranded) (Kim, Langmead et al. 2015), and estimated the read counts per gene by using htseq-count (v.0.9.1, union mode, reverse stranded) (Anders, Pyl et al. 2015) available on Galaxy Project server (https://usegalaxy.org/). Differential gene expression was analyzed by DESeq2 (v.2.11.40.6) (Love Mi., Huber W. et al. 2014) and genes with absolute Log2 fold change ≥ 1 and p.adjust < 0.05 were considered to be differentially expressed. Heatmaps and Gene ontology (GO) enrichment analysis were generated by R studio using the Bioconductor packages ComplexHeatmap (Gu, Eils et al. 2016) and clusterProfiler (Yu, Wang et al. 2012), respectively. Sequencing data were submitted to GEO Datasets under accession number GSE173292.

### Flow cytometry and cell cycle analysis

Cells were harvested with 0.05% Trypsin-EDTA to preserve the surface antigens and resuspended in Flow staining buffer (1% w/v BSA and 0.25 mM EDTA pH 8 in PBS) 100 μl per 100,000 cells. For detection of cell stemness, the samples were stained with TRA-1-60 conjugated FITC antibody (R&D system, FAB4770G, 1:50 dilution) or normal IgM conjugated FITC (Sigma, SAB4700699, dilution 1:50) for 30 min at 4°C in dark and washed once before acquisition. To determine early neural differentiation, mouse hybridoma antibody raised against glial precursor marker, A2B5 (DSHB, mAb 4D4) was used at dilution 1:20 and incubated with the cell solution at 4°C for 30 min before staining with AlexaFlour 488 goat anti-mouse (Cell Signaling, 1:100 dilution) for another 30 min at 4°C, in dark. Cells were analyzed with Accuri C6 plus Flow Cytometer (BD Biosciences) and data processed with FlowJo software. Unstained cells and isotype/negative controls from siCtrl were used for gating the cell population.

For cell cycle analysis, the harvested cells were resuspended in Vindelov’ reagent (10 mM Tris pH 7.4, 10 mM NaCl, 50 μg/mL Propidium iodide, 0.1% v/v NP40, 0.1 mg/mL RNaseA) and incubated for 1-2 h at 4°C in dark prior to analysis. The samples were run at slow rate (150 cells/sec or less) and the “bare nuclei” were first set the threshold by FSC and then gated using PI-Area vs. PI-Height to eliminate doublets. The percentage of cells in different phases was examined by FlowJo using integrated Cell Cycle platform.

### Statistical analysis

Comparisons between experimental conditions were done using GraphPad Prism software (GraphPad). The unpaired two-tailed Student’s t-test was used to analyze differences between two experimental groups. Data are shown as mean ± SD. The asterisks in each graph indicate statistically significant changes as *P < 0.05, ** P < 0.01, *** P < 0.001, **** P < 0.0001.

## Results

### Nuclear localization of DDX5 and DDX17 in hPSCs NTERA2 and their neural derivatives

To determine expression of DDX5 and DDX17 in PSCs and during their neural differentiation, real-time PCR, western blot, and immunofluorescence staining were performed using NTERA2 and their retinoic acid-induced neural derivatives. We confirmed that expression of *OCT4* and *NANOG* is down-regulated, while that of *PAX6, NESTIN, SOX1*, and *TUBB3* is up-regulated suggesting neural differentiation of the stem cells upon the induction of retinoic acid (Supplementary Figure S1). In contrast to those stem cell and differentiation markers, expression of DDX5 and DDX17 is relatively maintained during the course of differentiation (Figures 1A-1B). Immunofluorescent staining showed an overlapping pattern of DDX5 or DDX17 with DAPI in both PSCs and their neural derivatives (Figure 1C). As expected, DDX5 and DDX17 are expressed in cells positive for OCT4 and TUBB3, which represent PSCs and neurons, respectively. This result indicates that the orthologous RNA helicases DDX5 and DDX17 are expressed throughout neural differentiation, and preferentially localize in the nucleus.

**Figure 1.**
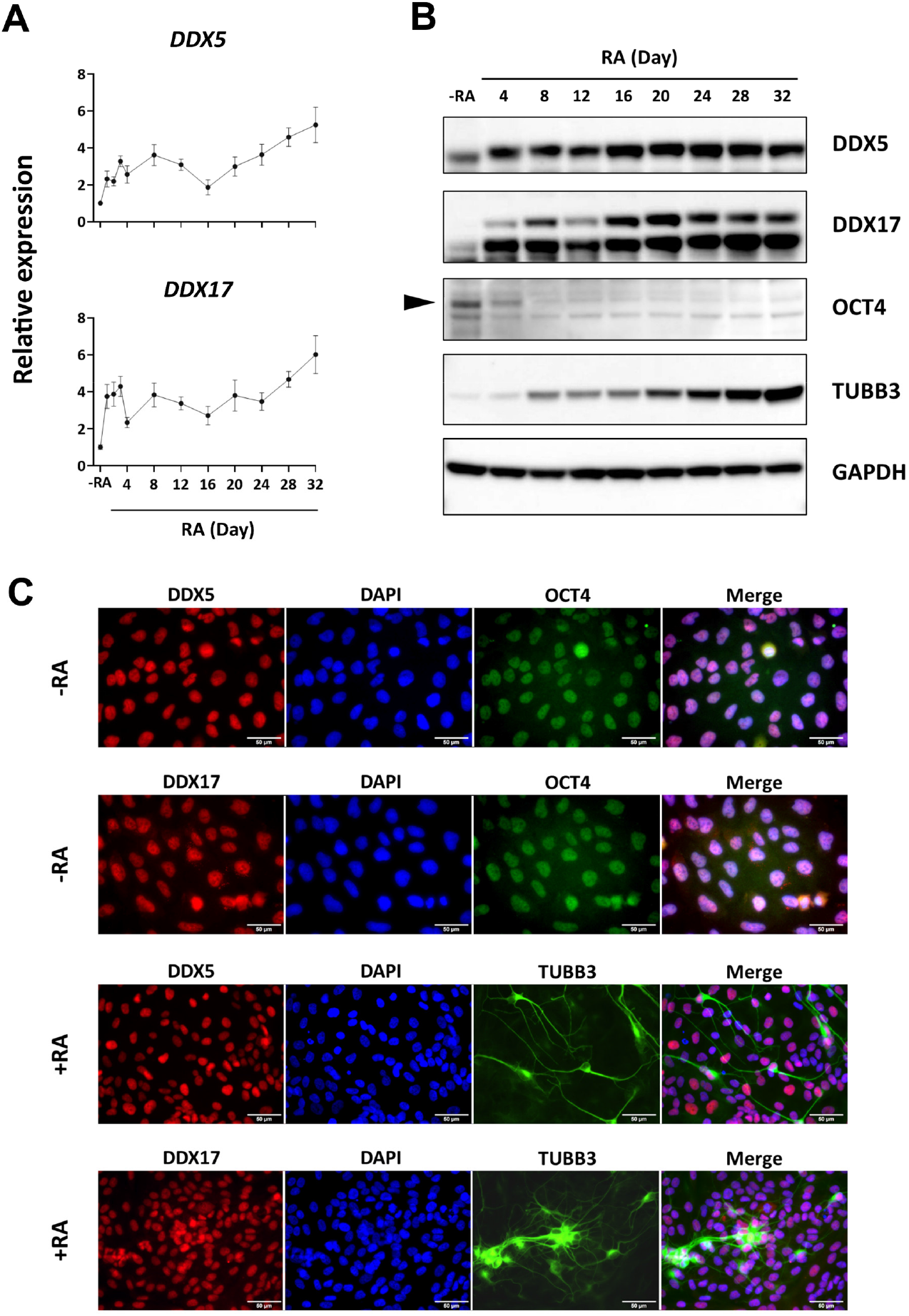
The RNA helicases DDX5 and DDX17 are expressed during neural differentiation, and preferentially localize in the nucleus. NTERA2 cells were seeded at 10,000 cell/cm^2^ and were induced to differentiate toward neural lineage by retinoic acid (RA) at 10 μM for the time course of 32 days. Samples were collected every 4 days for real-time PCR (A). *GAPDH* was utilized as an internal control. Error bars represent SD; (n=3). (B) Western blots of DDX5, DDX17, the stem cell marker OCT4, and the neural differentiation marker TUBB3. GAPDH was utilized as an internal control. (C) Immunofluorescent staining of undifferentiated culture and day 32 retinoic acid-induced neural derivatives was performed using anti-OCT4 and anti-TUBB3 antibodies (green), respectively. The cells were co-stained with anti-DDX5 or anti-DDX17 (red). DAPI was used to localize the nucleus. Scale bar = 50 μm.

### Genome-wide localization of DDX5 and DDX17 in hPSCs NTERA2 and their neural derivatives

Due to their nuclear localization in the stem cells and their neural counterparts, we next established the role of DDX5 and DDX17 in genome regulation of the cells. Chromatin immunoprecipitation and sequencing (ChIP-seq) of the RNA helicases was performed for hPSC and retinoic acid-induced neural cultures differentiated for 14 days. We have previously reported a total of 14,131 genome-wide binding sites of DDX5 in NTERA2 (Wongtrakoongate, Riddick et al. 2015). ChIP-seq of the DDX5 orthologs DDX17 in the stem cells further identified 6,058 binding sites genome-wide. In the neural derivatives, we found 13,158 and 7,673 genomic binding sites of DDX5 and DDX17, respectively (Supplementary Table S1). Among these binding sites, 647 and 884 binding sites are over-lapped by both DDX5 and DDX17 in hPSCs and their neural derivatives, respectively (Figure 2A). By analyzing the genomic distribution of the RNA helicases, approximately 25.0% and 14.5% of total DDX5 and DDX17 binding sites in hPSCs could be categorized as the promoter regions. However, when compared with the stem cells, differentiated neural cultures possess a decrease in promoter occupancy, with 12.8% and 4.0% of total DDX5 and DDX17 promoter binding sites, respectively (Figure 2B). In contrast, chromatin of the neural derivatives harbors higher proportions of distal intergenic region-associated DDX5 and DDX17 occupancy sites than the stem cells, with 45.1% and 54.2% in the differentiated cells and 35.8% and 42.9% in the stem cells, respectively. These distal intergenic regions might possess an enhancer function as described previously (Van Bakel, Nislow et al. 2010, Djebali, Davis et al. 2012). Gene ontology (GO) analysis uncovered the chromatin occupancy of DDX5 and DDX17 at genes involved in embryonic development, organogenesis, morphogenesis, neural development, and neural function (Figure 2C). These ChIP-seq analyses indicate the genome-wide binding of the two RNA helicase orthologs at differentiation-associated genes, with a minority of their binding sites being co-occupied by both DDX5 and DDX17.

**Figure 2.**
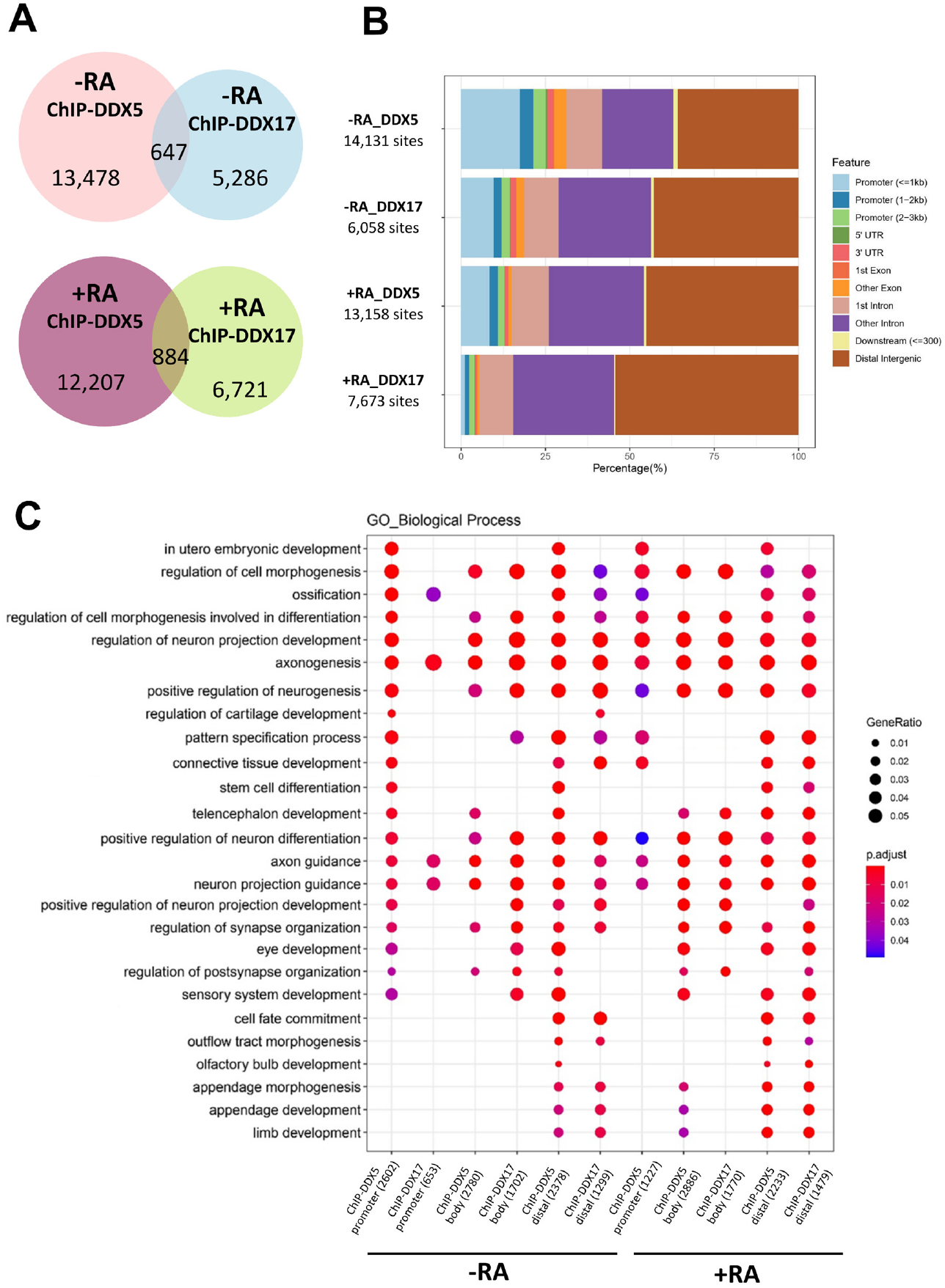
DDX5 and DDX17 occupy the chromatin genome-wide, but a minority of DDX5- and DDX17-occupied genomic regions are co-occupied by both RNA helicases. ChIP-seq of DDX5 and DDX17 was performed using the hPSCs NTERA2 and their neural derivatives prepared from retinoic acid treatment for 14 days. (A) Venn diagrams of regions bound by DDX5 and DDX17 in NTERA2 (top) and the neural derivatives (bottom) were generated by the library (VennDiagram). (B) Genomic occupancy of DDX5 and DDX17 both in undifferentiated and retinoic acid-induced cultures was analyzed using ChIPseeker to assign genetic elements. (C) Gene ontology related to differentiation processes was analyzed by clusterProfiler and grouped according to genetic elements including promoters, gene bodies, and distal intergenic regions. The number in parentheses refers to number of genes associated with each category.

### Genome-wide transcriptional control of DDX5 and DDX17 in hPSCs NTERA2 and their neural derivatives

To elucidate whether the orthologs DDX5 and DDX17 control gene expression transcriptome-wide, expression of DDX5 and DDX17 was depleted by RNA silencing using ON-TARGETplus SMARTpool siRNA. Real-time PCR, western blot, and immunofluorescence staining confirmed a reduction of DDX5 and DDX17 expression (Supplementary Figures S2A-S2C). Upon silencing of DDX5 and/or DDX17, RNA sequencing (RNA-seq) was performed for hPSC and retinoic acid-induced neural cultures differentiated for 14 days. RNA-seq reads were mapped to the genome reference hg19/GRCh37. Analysis of differentially expressed genes (DEGs) was performed with DESeq2 under the Galaxy Project. By using the parameters of absolute Log2 fold change ≥ 1 and p.adjust < 0.05, the silencing of *DDX5* and *DDX17* transcripts was confirmed in all samples (Figure S3). However, we could only retrieve 0 and 20 genes whose transcripts are differentially expressed in hPSCs individually silenced for DDX5 and DDX17, respectively. Similarly, we retrieved 0 and 46 genes whose transcripts are differentially expressed in the neural derivatives individually silenced for DDX5 and DDX17, respectively. In contrast, by simultaneous silencing of both DDX5 and DDX17, we retrieved 2,071 and 1,132 genes whose transcripts are differentially expressed in hPSCs and their neural derivatives, respectively (Figures 3A-3B and Supplementary Table S2). By using DESeq2 and ComplexHeatmap, we generated the heapmaps of DEGs with their Log2 fold change for hPSCs (Figure 3C) and their neural derivatives (Figure 3D). Examples of genes whose function is involved with neuroectodermal differentiation and organogenesis are indicated. Due to the few DEGs of the single knockdowns, we were unable to perform GO analysis of these samples. Nevertheless, GO analysis using DEGs of hPSCs simultaneously depleted for DDX5 and DDX17 showed that the GO groups of neural development such as axonogenesis are enriched for genes down-regulated by DDX5/DDX17 knockdown (Figure 3E). On the other hand, the GO groups of multilineage developments such as differentiation of mesoderm lineages and trophoblasts are enriched for genes up-regulated by DDX5/DDX17 knockdown in hPSCs. In comparison to hPSCs, silencing of both DDX5 and DDX17 in the neural derivatives led to DEGs enriched for non-neuroectodermal lineages apart from the GO group of gliogenesis. These results suggest that both DDX5 and DDX17, but not individuals, mutually control the transcriptional expression of differentiation-associated genes in hPSCs and their neural derivatives, and, in particular, facilitate the expression of neurogenesis-related genes in hPSCs.

**Figure 3.**
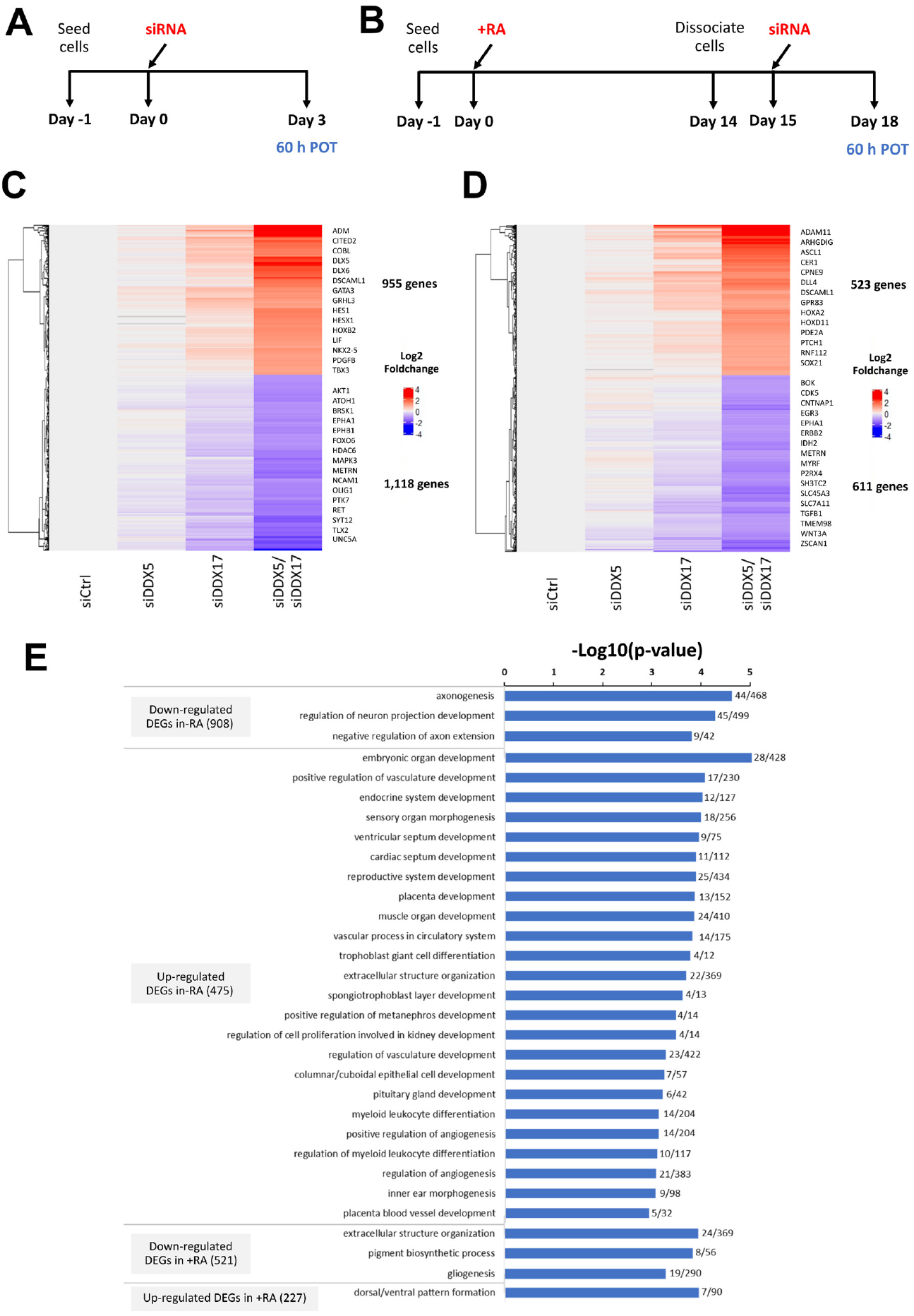
DDX5 and DDX17 facilitate transcriptional expression of neurogenesis-related genes in hPSCs. Experimental outline. The culture of undifferentiated (A) or NTERA2-induced neural differentiation by retinoic acid at day 14 (B) was pre-treated for 60 h with control siRNA or siDDX5/DDX17 and determined the knockdown efficiency and RNA quality prior to preparation of RNA-seq library by real-time PCR and bioanalyzer, respectively. The RNA helicases DDX5 and DDX17 control expression of 2,071 and 1,132 genes in the hPSCs NTERA2 and their neural derivatives, respectively. RNA-seq was performed for cells depleted for either DDX5, DDX17, or both using ON-TARGETplus SMARTpool siRNA. Following silencing of both DDX5 and DDX17 in hPSCs, 1,116 and 955 genes are down- and up-regulated in DDX5/DDX17 knockdown cells, respectively. For the neural derivatives, 609 and 523 genes are down- and up-regulated in DDX5/DDX17 knockdown cells, respectively. Heat maps of DEGs between DDX5/DDX17 double knockdowns and the controls were analyzed for hPSCs (C) and the neural derivatives (D) using DESeq2 and ComplexHeatmap. Examples of genes associated with neural development or embryogenesis are shown for each heatmap. (E) DEGs regulated by both DDX5 and DDX17 in hPSCs or the neural derivatives were analyzed for categories of enriched gene ontologies (GO) using clusterProfiler. The number shown for each bar chart represents the ratio of identified genes to all genes within a given GO.

### DDX5 and DDX17 do not maintain stemness of hPSCs

Next, we aimed to ascertain whether the RNA helicases DDX5 and DDX17 control pluripotency and neural differentiation. The stem cell surface marker TRA-1-60, which is a pristine marker for the pluripotent state (Adewumi, Aflatoonian et al. 2007, Chan, Ratanasirintrawoot et al. 2009), was employed to determine the stem cell state of hPSC and retinoic acid treated sample (Figure 4A). Using flow cytometry, we did not observe changes in the number of cells positive for TRA-1-60 in PSC cultures (Figures 4B-4C). However, silencing of both *DDX5* and *DDX17*, but not individuals, led to an increase in the expression level of TRA-1-60 (Figure 4D). Upon treatment of retinoic acid for 3 days, the numbers of TRA-1-60-positive cells remain higher in cells individually depleted for DDX5 and DDX17 as well as cells with double knockdown than that of the control suggesting that the RNA helicases might facilitate the exit of pluripotent state (Figures 4E-4G). To test whether cell cycle profile is altered by silencing of the RNA helicases, we performed propidium iodide staining followed by flow cytometry. We found that the depletion of both RNA helicases in hPSCs resulted in an increase in G2/M fraction at the expense of G1 and S fractions (Figure 4H). Nonetheless, we observed no changes in transcriptional expression of the stem cell markers *OCT4, NANOG*, and *SOX2* (Figure 4I). These results suggest that the RNA helicases DDX5 and DDX17 do not maintain the stem cell state, but, rather, facilitate retinoic acid-induced differentiation of hPSCs.

**Figure 4.**
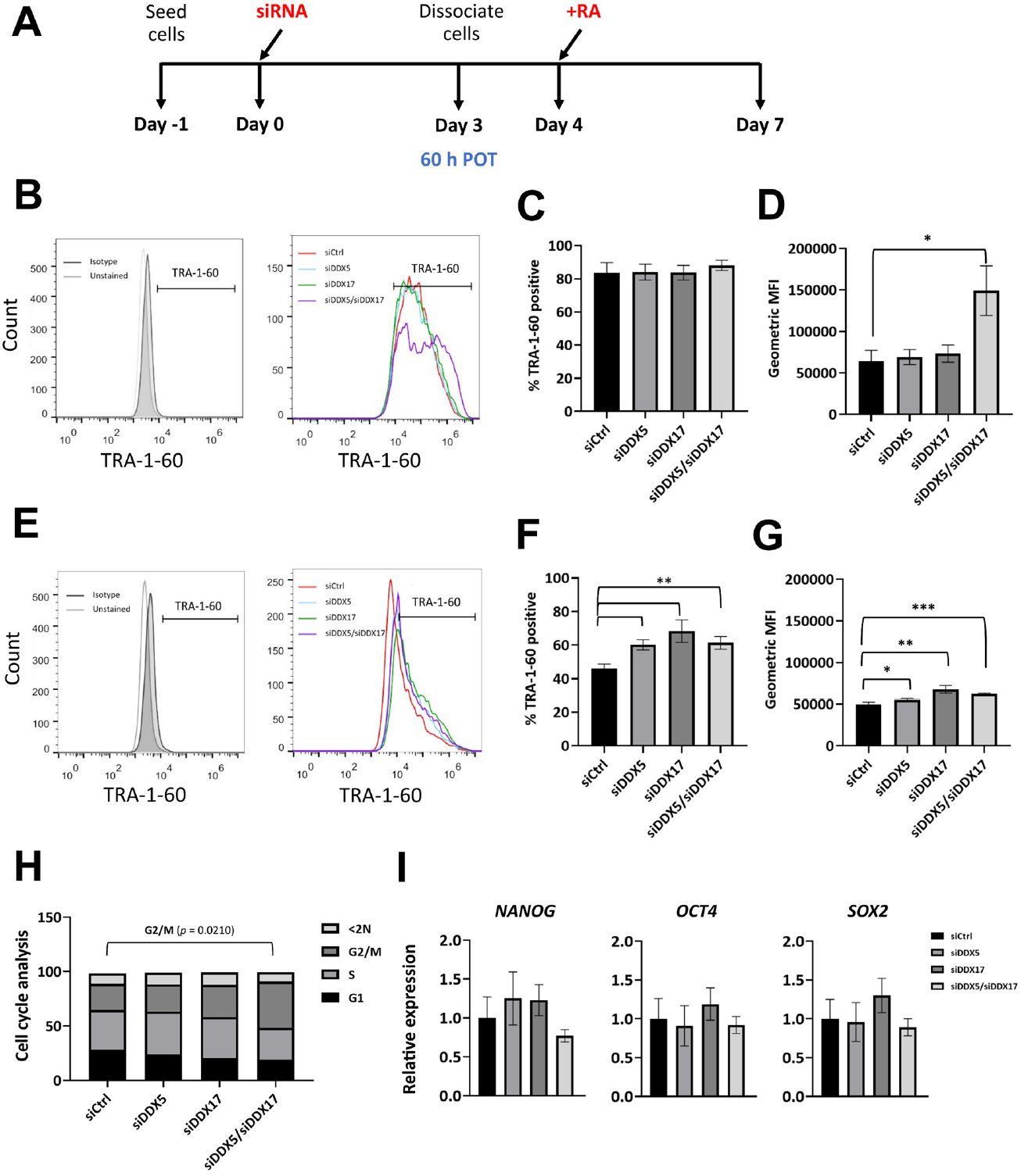
The RNA helicases DDX5 and DDX17 do not maintain the stem cell state of NTERA2. Experimental outline for figure 4B-4G is shown in (A). The hPSC culture was silenced for individual RNA helicases or both using siRNA for 60 h and then dissociated to be analyzed by flow cytometry or further induced differentiation by retinoic acid for 3 days. Flow cytometric analyses of the stem cell marker TRA-1-60 in the hPSCs NTERA2 after 60-hour post-transfection (POT) (B-D) and day 3 retinoic acid-induced cells (E-G) show that depletion of both DDX5 and DDX17 attenuated differentiation of the stem cells as determined by the decrease in cells positive for TRA-1-60. The TRA-1-60 histograms are shown to compare the percentage of positive populations and expression level (geometric mean fluorescent intensity; MFI). (H) Cell cycle analysis was performed using propidium iodide (PI) and was represented according to the fraction of DNA content. (I) The relative gene expression level of stem cell markers in the hPSCs NTERA2 after 60-hour POT was determined by real-time PCR. *GAPDH* was utilized as an internal control. Error bars represent SD; (n=3). Student t-test. * p<0.05; ** p<0.01; *** p<0.001.

### DDX5 and DDX17 are both mutually important for early neural differentiation of hPSCs

Since the RNA helicases, DDX5 and DDX17 are expressed in hPSCs and during neural differentiation (Figure 1) and control transcriptional expression of neurogenesis-related genes (Figure 3E), we asked whether they regulate neural differentiation of hPSCs by performing the RNAi knockdown in the PSC culture. To determine early neural differentiation, we employed the cell surface marker A2B5, which demarcates early neural precursor cells (Adewumi, Aflatoonian et al. 2007). In comparison with the hPSCs NTERA2 induced by retinoic acid for 8 days, which showed approximately 80% positive for A2B5, cells depleted for both DDX5 and DDX17, but not individuals, possess only 60% of A2B5-positive cells (Figures 5A-5D). Using a neurosphere induction comprising of N21, bFGF, and EGF, we found that the number of neurospheres, but not their size, is reduced by depletion of both DDX5 and DDX17, but not individuals (Figures 5E-5G). Additionally, real-time PCR analyses performed using retinoic acid-induced hPSCs in a time-course manner of 32 days showed that transcriptional expression of neurogenic transcription factors including *SOX2, SOX21, SOX1, PAX6, ASCL1*, and *NEUROG2* is down-regulated throughout the differentiation in cells silenced for both DDX5 and DDX17 but not individuals (Figure 6A-6B). On the other hand, silencing of *DDX5* and *DDX17* in neural derivatives derived from PSCs induced by retinoic acid for two weeks did not alter expression of the neurogenic transcription factors, except for *PAX6* (Figure 6C-6D), suggesting that DDX5 and DDX17 do not regulate expression of these genes in the neural derivatives. These results indicate a key role of both RNA helicases DDX5 and DDX17 in the early neural differentiation of hPSCs.

**Figure 5.**
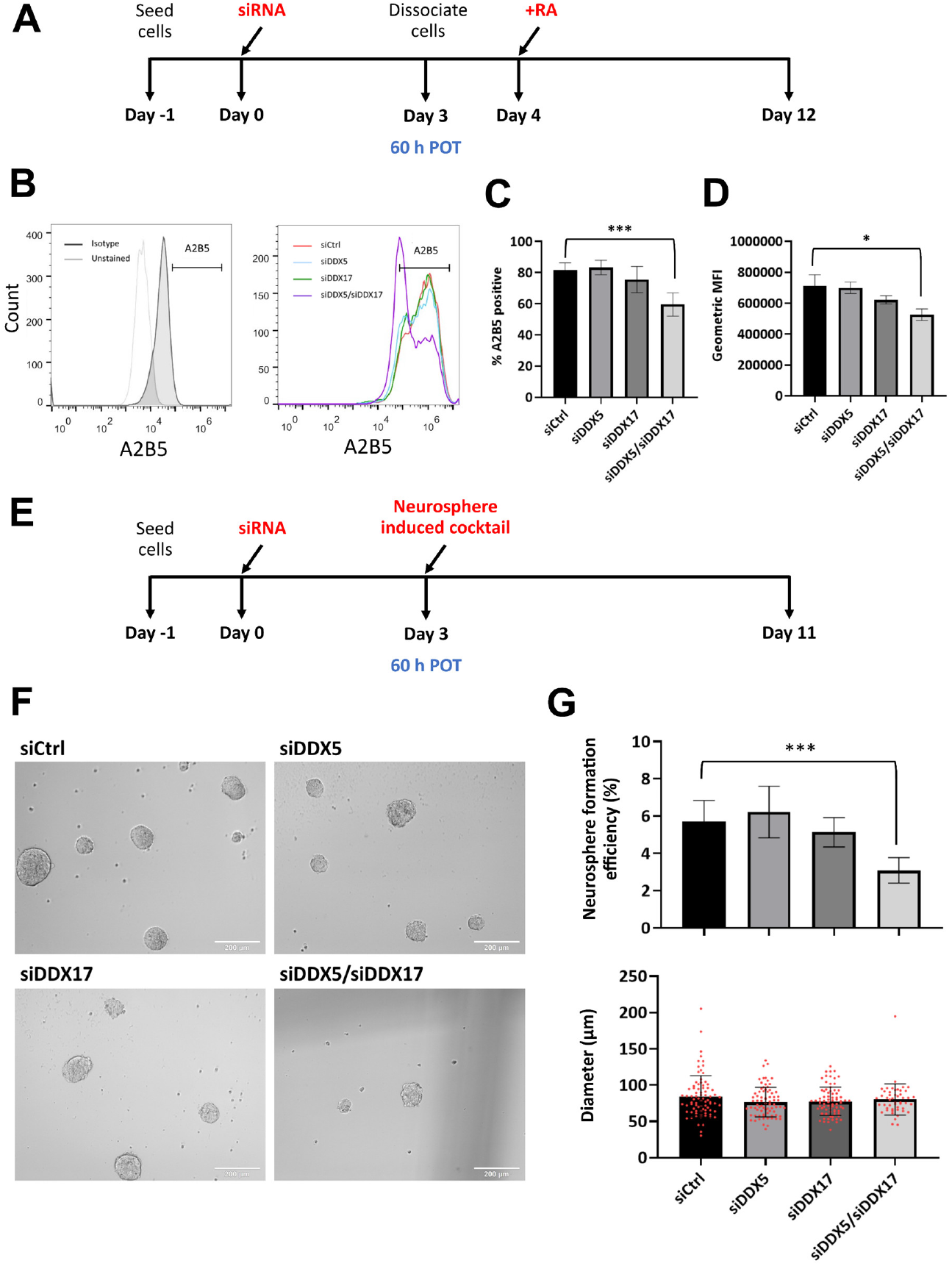
The RNA helicases DDX5 and DDX17 facilitate early neural differentiation of hPSCs. Experimental outline. The undifferentiated NTERA2 cells were transfected with control siRNA, siDDX5, siDDX17, or siDDX5/siDDX17 for 60 h prior to re-seeding in growth medium containing retinoic acid (A) or N21, bFGF, and EGF cocktail to induce neurosphere formation (E). Flow cytometric analyses of the early neural differentiation marker A2B5 in the hPSCs NTERA2 show that depletion of both DDX5 and DDX17 attenuated neural differentiation of the stem cells as determined by the increase in cells positive for A2B5 (B-D). The A2B5 histograms are shown to compare the percentage of positive populations and expression level (geometric mean fluorescent intensity; MFI). (F) Representative images of neurospheres induced from hPSCs NTERA2 after 8 days. Scale bar = 200 μm. (G) Silencing of both DDX5 and DDX17 reduced the efficiency of neurosphere formation (Top), whereas the size (in diameter) of the neurospheres is not altered (bottom). Student t-test. * p<0.05; ** p<0.01; *** p<0.001.

**Figure 6.**
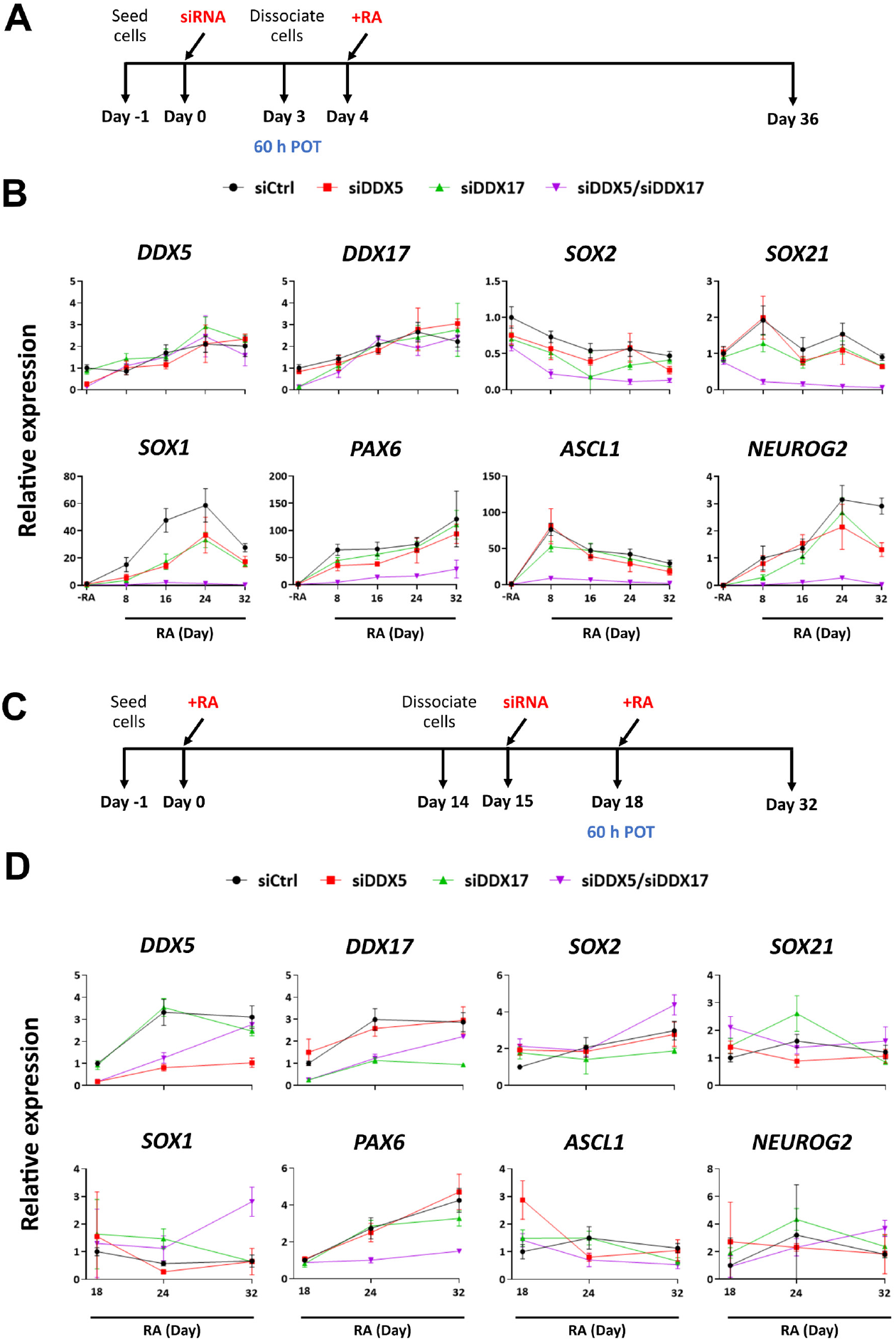
Early suppression of DDX5/DDX17 in the pluripotent cell stage sustainably reduces the expression of neural-specific genes. Experimental outline. The culture of undifferentiated (A) or NTERA2-induced neural differentiation by retinoic acid at day 14 (C) was pre-treated for 60 h with control siRNA or siDDX5/DDX17 prior to induced differentiation by retinoic acid until reaching the mature neuron cell stage. (B and D) The relative gene expression level of key neurogenic transcription factors was determined by real-time PCR upon retinoic induction for the time course of 32 days. *GAPDH* was utilized as an internal control. Error bars represent SD; (n=3).

Next, to elucidate whether DDX5 and DDX17 facilitate neurogenesis of neural progenitors and neurons from hPSCs, we performed immunofluorescence staining of cells expressing NESTIN and TUBB3, which determine progenitor and mature states of neural derivatives, respectively. Expression of the two RNA helicases was individually or simultaneously silenced in the hPSCs NTERA2. The cells were then treated with retinoic acid for 16 days to induce neural differentiation.

Silencing of either DDX5 or DDX17 did not affect the numbers of NESTIN- or TUBB3-positive cells. On contrary, hPSCs silenced for both RNA helicases generate fewer NESTIN- or TUBB3-positive cells compared with the control (Figures 7A, 7C-7D). In order to test whether the role of the RNA helicases in neurogenesis can be seen in neural derivatives induced from hPSCs, NTERA2 cells were induced to differentiate for 14 days before silencing of the RNA helicases. The neural derivatives depleted for *DDX5* or *DDX17* or both were then induced for further differentiation until day 32. Nonetheless, we did not observe changes in the numbers of NESTIN- or TUBB3-positive in cells induced from the neural derivatives (Figures 7B, 7C-7D), indicating that the two RNA helicases do not control neurogenesis at this later stage of neural development. Together, these results suggest that both RNA helicases DDX5 and DDX17 mutually facilitate the generation of neural progenitors and mature neurons from hPSCs but not from neural derivatives induced from the stem cells.

**Figure 7.**
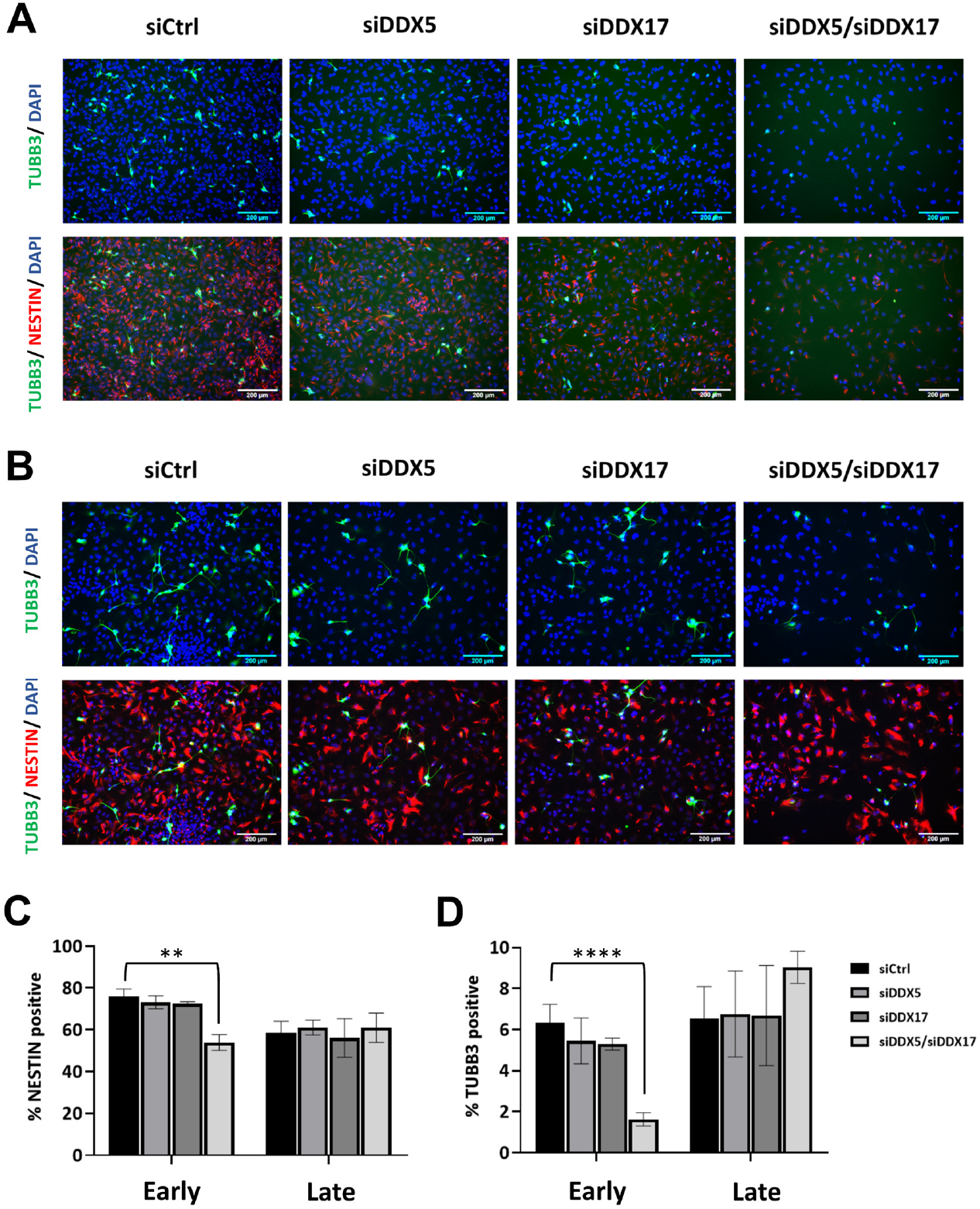
Both RNA helicases DDX5 and DDX17 mutually facilitate the generation of neural progenitors and mature neurons from hPSCs. (A) The stem cells were silenced for individual RNA helicases or both using siRNA and were treated by retinoic acid for 16 days to derive early neural derivatives. (B) hPSCs were induced by retinoic acid to differentiate for 14 days to obtain early neural derivatives prior to RNA silencing. Upon knockdown of the RNA helicases, the neural derivatives were allowed to differentiate further in the presence of retinoic acid for another 14 days to derive late neural derivatives. Immunofluorescent staining of mature neurons and neural progenitors was performed using anti-TUBB3 (green) and anti-NESTIN (red) antibodies, respectively. The cells were counter stained with DAPI. Scale bar = 200 μm. Fluorescence micrographs of the early neural derivatives (A) and the late neural derivatives (B) were determined for the percentage of cells positive for NESTIN (C) or TUBB3 (D). Error bars represent SD; (n=3). Student t-test. ** p<0.01; **** p<0.0001.

## Discussion

The enzymatic and molecular mechanisms underlying the two RNA helicase orthologs DDX5 and DDX17 have been widely characterized. However, little is known for their phenotypic regulation in human neurogenesis. In the present study, we have determined that the two RNA helicases DDX5 and DDX17 facilitate neural differentiation of the hPSCs NTERA2.

We find that expression of DDX5 and DDX17 is increased in retinoic acid-induced neural derivatives compared to their undifferentiated counterparts, even though all cells essentially express the two RNA helicases (Figure 1). Though there is no report on the expression of DDX5 during human embryonic development, expression of DDX5 protein has been reported to be developmentally up-regulated in various tissues of mouse fetuses (Stevenson, Hamilton et al. 1998). In cell line models of human neurogenesis, *DDX17* transcript was identified by RNA fingerprinting by arbitrarily primed PCR (RAP-PCR) to be down-regulated during retinoic acid-induced differentiation of the hPSCs NTERA2 (Cheung, Chu et al. 1997). In the human neuronal progenitor cell lines, SH-SY5Y and CLBMA2, DDX5 and DDX17 proteins are down-regulated by retinoic acid treatment (Lambert, Terrone et al. 2018). In contrast, another study has shown that expression of *DDX5* and *DDX17* transcripts are unperturbed by treatment of retinoic acid in the human neuronal progenitor cell line SH-SY5Y (Ip, Chung et al. 2000). These and our studies suggest cell type- and lineage-specific expression patterns of the two RNA helicases. We also confirm the nuclear localization of DDX5 and DDX17 in both PSCs and their neural derivatives. This nuclear localization profile is also consistent with previous reports on DDX5 and DDX17 (Lamm, Nicol et al. 1996, Kahlina, Goren et al. 2004, Alqahtani, Gopal et al. 2016, Weise, Arumugam et al. 2019), whereas other RNA helicases such as DDX3 (Sekiguchi, Iida et al. 2004, Vakilian, Mirzaei et al. 2015) and DDX6 (Smillie and Sommerville 2002, Kami, Kitani et al. 2018) have been identified to mostly localize in the cytoplasm. It is yet to be determined whether DDX5 and DDX17 can possess functions in the nucleus apart from being transcriptional coregulators (Dardenne, Polay Espinoza et al. 2014, Giraud, Terrone et al. 2018) and microRNA processors (Hong, Noh et al. 2013, Kao, Cheng et al. 2019, Weise, Arumugam et al. 2019).

We have previously reported a genome-wide occupancy of DDX5 in the hPSCs NTERA2, which reveals genomic binding sites associated with embryonic- and differentiation-related genes (Wongtrakoongate, Riddick et al. 2015). In the current report, we have employed NTERA2 cells and their neural derivatives derived from retinoic acid induction to characterize the genome-wide occupancy of DDX5 and DDX17 (Figure 2). Even though the two RNA helicases have been shown to form a heterodimer in vitro and in vivo (Ogilvie, Wilson et al. 2003, Mersaoui, Yu et al. 2019), only a minority of DDX5- and DDX17-binding sites are overlapped (Figure 2). Several findings have been proposed for delivery mechanisms of DDX5 or DDX17 at target promoters. For example, DDX5 has been shown to be delivered by the transcription factors CTCF (Yao, Brick et al. 2010), Fra-1 (He, Song et al. 2019), NANOG (Wongtrakoongate, Riddick et al. 2015), respectively. Similarly, DDX17 has been reported to be delivered by REST (Lambert, Terrone et al. 2018) and SOX2 (Alqahtani, Gopal et al. 2016). Nonetheless, both DDX5 and DDX17 can also be co-delivered by MyoD and Runx2 in myogenic cells (Caretti, Schiltz et al. 2006) and osteogenic cells (Jensen, Niu et al. 2008), respectively. Thus, DDX5 and DDX17 might independently form complexes with different transcription factors, leading to differential chromatin occupancy. ChIP-seq of these two orthologs in standard cell lines/types should further facilitate understanding how DDX5 and DDX17 are deposited to chromatin.

DDX5 has been individually shown for its transcriptome-wide gene regulation (He, Song et al. 2019, Legrand, Chan et al. 2019). Here, using the hPSCs NTERA2 and their neural derivatives as the models, we did not observe differential transcriptome-wide alteration of either DDX5 or DDX17 knockdown cells (Figure 3). Rather, when being simultaneously silenced, more than thousands of differentially expressed genes can be identified in both NTERA2 and the neural derivatives. Function of these genes is associated with neural differentiation and embryonic development, supporting the chromatin occupancy of DDX5 and DDX17 at genes related to these processes. Similar findings of the synergistic role of DDX5 and DDX17 have been reported for cell proliferation and rRNA processing (Jalal, Uhlmann-schiffler et al. 2007) and NFAT5-dependent cancer invasion (Germann, Gratadou et al. 2012). With regard to the pluripotent state, genes differentially expressed in DDX5/DDX17 double knockdown cells are not involved in the maintenance of stem cell state. In addition, expression of *NANOG, OCT4*, and *SOX2* is unperturbed in the double knockdown, consistent with the relatively constant numbers of cells positive for TRA-1-60 (Figure 4). Yet, we find that both DDX5 and DDX17 are important for early differentiation, since the number of TRA-1-60 positive cells of the control knockdown under retinoic acid treatment was lower than that of DDX5/DDX17 knockdown. Our finding therefore supports the role of DDX5, and possibly DDX17, as the barrier of pluripotency (Li, Lai et al. 2017, Li, Song et al. 2020).

As being the multifunctional RNA helicases, DDX5 and DDX17 have been extensively elucidated for their transcriptional involvement in various cellular contexts including developmental processes (Fuller-Pace 2013). In the present study, we report that the two RNA helicases are critical for neural differentiation of hPSCs (Figure 5 and 6), possibly due to their contribution to up-regulation of neurogenic transcription factors including *SOX2, SOX21, SOX1, PAX6, ASCL1*, and *NEUROG2*. DDX5 and DDX17 have been shown to promote neuronal differentiation of the human neuroblastoma cell line SH-SY5Y through both REST-mediated transcriptional and post-transcriptional controls (Lambert, Terrone et al. 2018), while they facilitate self-renewal of mouse neural progenitors through destabilization of differentiation-related mRNAs (Moon, Bai et al. 2018). Apart from neurogenesis, DDX5 and/or DDX17 have been shown to promote myoblast and osteoblast differentiation via interaction with the transcription factors MyoD (Caretti, Schiltz et al. 2006) and Runx2 (Jensen, Niu et al. 2008), respectively. In conclusion, the present study uncovers the transcriptional regulatory role of DDX5 and DDX17 in the neural differentiation of hPSCs. This finding provides novel insights into our understanding of RNA helicases underlying early embryonic neurogenesis.

## Acknowledgements

This research project was supported by CIF Grant, Faculty of Science, Mahidol University. PS was supported by Science Achievement Scholarship of Thailand. P.W. was supported by Mahidol University (New Discovery and Frontier Research Grant; grant number NDFR 11/2563) and the Office of National Higher Education Science Research and Innovation Policy Council by Program Management Unit for Human Resources and Institutional Development, Research and Innovation (PMU-B; grant number B05F630081). GF was supported by the Intramural Research Program of the National Institute of Diabetes and Digestive and Kidney Diseases, NIH. We are indebted to PW lab members for their suggestions and comments.

## Data availability

The data discussed in this publication have been deposited in NCBI’s Gene Expression Omnibus (GEO) database under accession code GSE174062 (including the SubSeries codes: GSE173292 and GSE174051) and DDX5 ChIP-seq data performing in wild-type NTERA2 cells has been previously deposited in GEO under accession code GSE58641.

---

**Supplementary S1.**
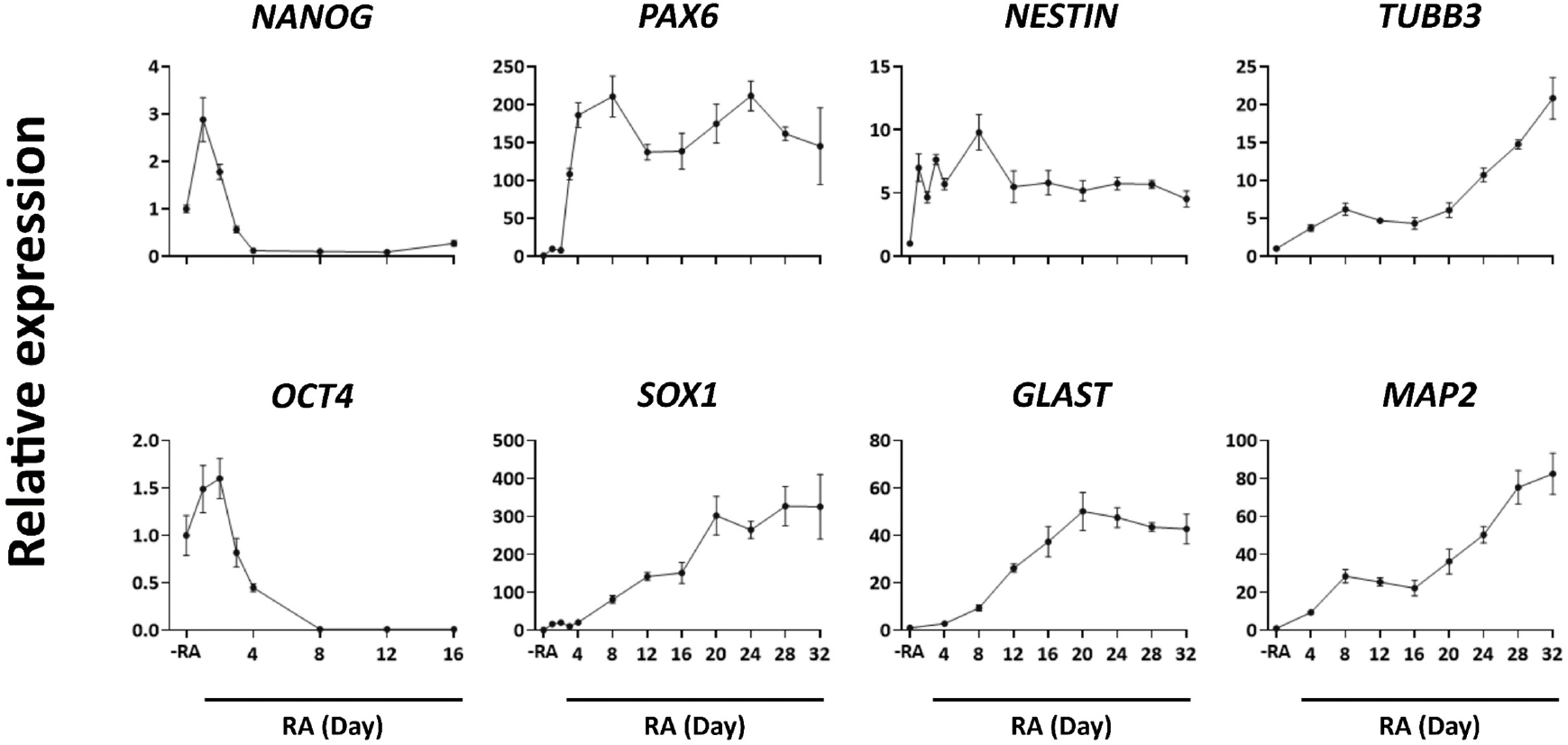
Neural differentiation of the hPSCs NTERA2 induced by retinoic acid treatment. NTERA2 cells were seeded at a density of 10,000 cell/cm^2^ and induced differentiation by 10 μM retinoic acid for 32 days. The cultures were collected with a 4-days interval and analyzed by real-time PCR. *NANOG* and *OCT4* were used as stem cell markers while *PAX6, NESTIN, SOX1, GLAST, TUBB3*, and *MAP2* were used as neural-specific markers. *GAPDH* was utilized as an internal control. Error bars represent SD; (n=3).

**Supplementary S2.**
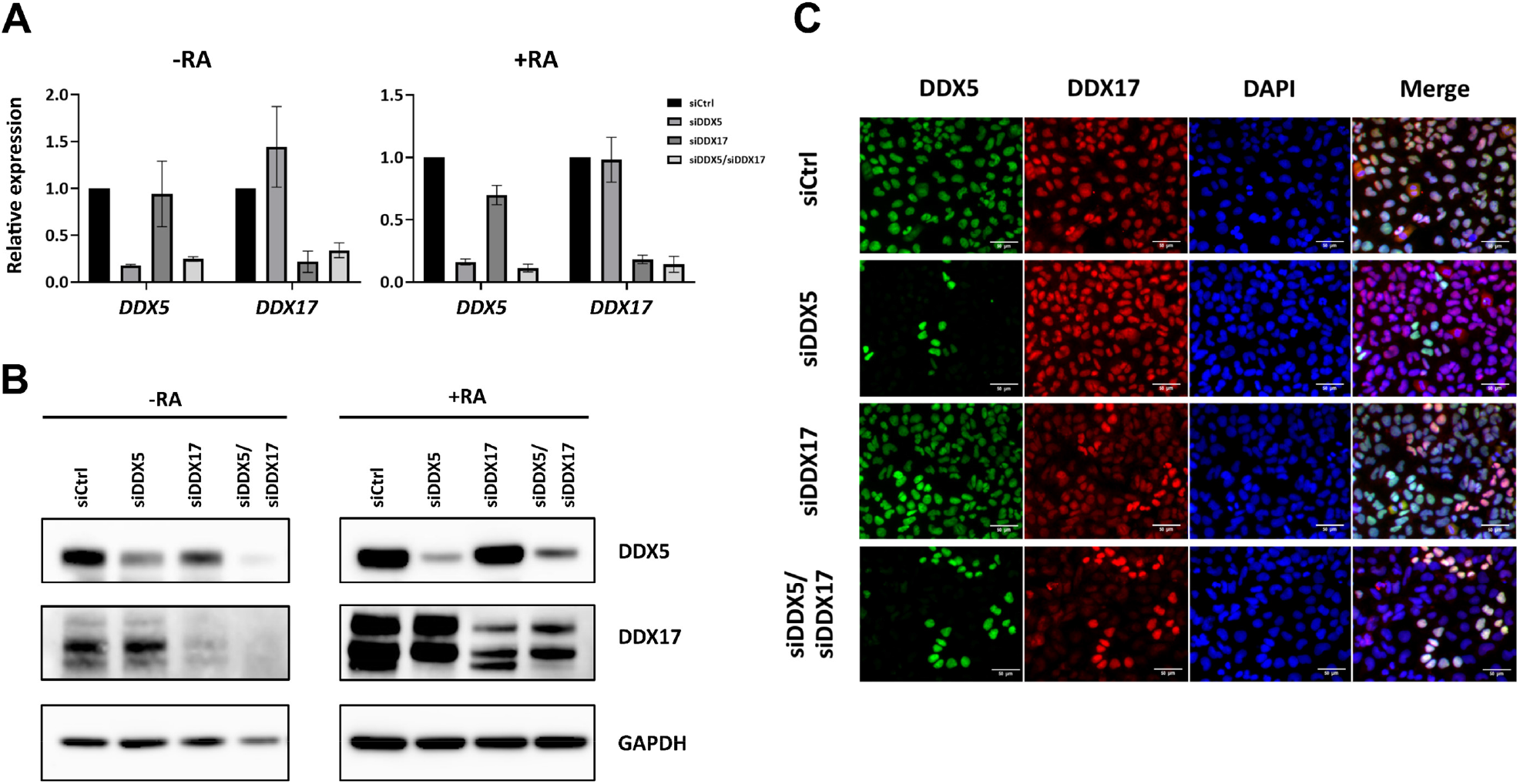
Silencing of DDX5 and DDX17. The culture of hPSCs and neural derivatives were transfected with an siRNA pool at concentration 10 μM each, and incubated for 60 hours prior to determination of transcript (A) and protein levels (B). The non-targeting siRNA was used as a control knockdown and GAPDH was utilized as an internal control for both real-time PCR and Western blot. Error bars represent SD; (n=3). (C) Simultaneous reduction of DDX5 and DDX17 double knockdown was confirmed by immunofluorescent staining of hPSC culture after treatment with either siDDX5, siDDX17, or both. The cells were co-stained with anti-DDX5 (green) and anti-DDX17 (red). DAPI was used to localize the nucleus. Scale bar = 50 μm.

**Supplementary S3.**
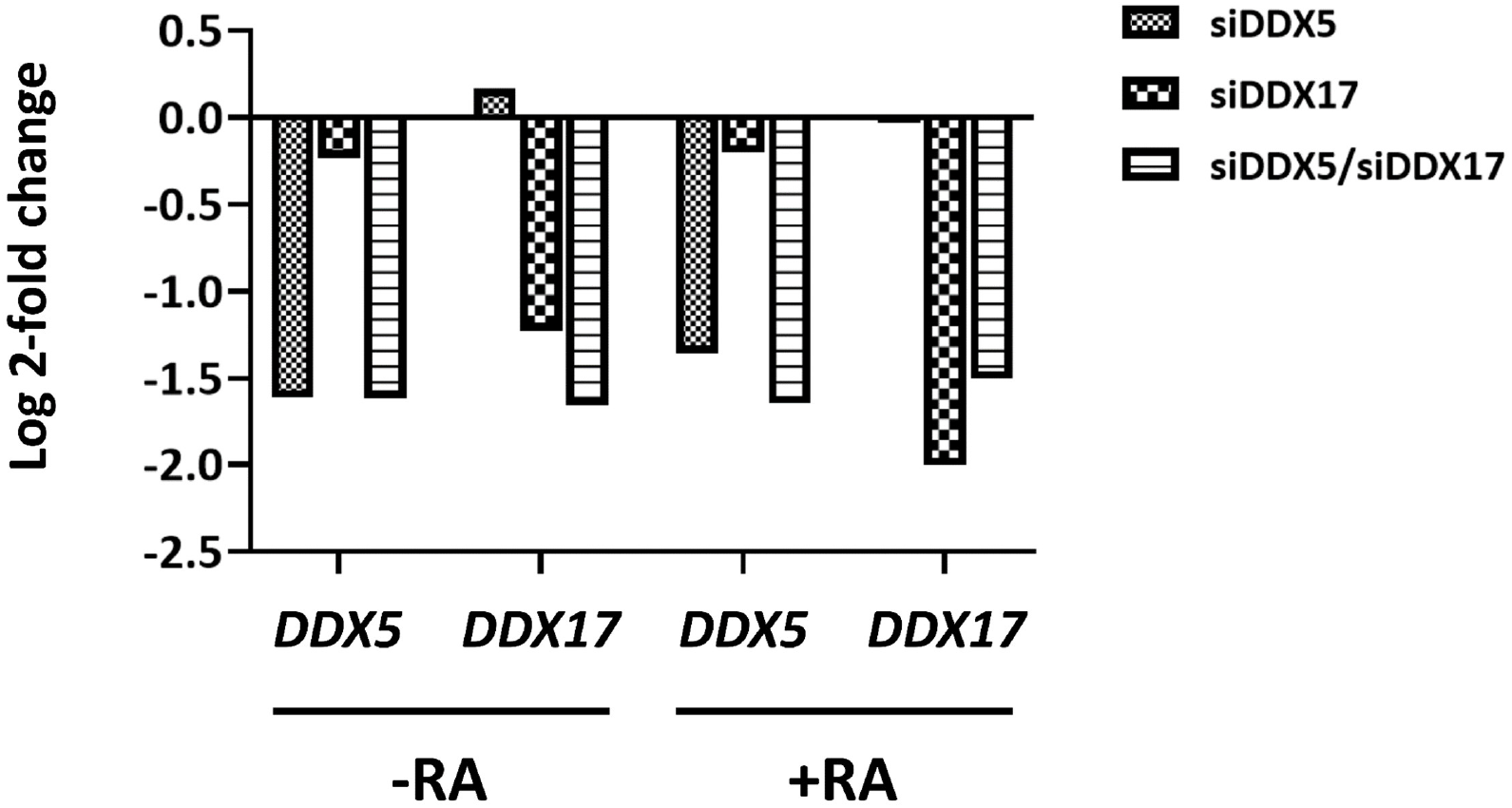
RNA-seq confirmed down-regulation *DDX5* and *DDX17*. The estimated counts of *DDX5* and *DDX17* transcripts were provided by htseq-counts and used as the input for differential expression analysis by DESeq2. The expression of DDX5 and DDX17 in single or double target silencing was calculated relative to non-target knockdown sample for both hPSC and the neural derivatives cultures and reported as Log2 fold change value. The statistical significance of DDX5 and DDX17 in all samples tested were observed with p.adjust < 0.0001.

